# Predicting RNA:DNA Triplex Structures from Sequence Features Using Deep Learning Architecture

**DOI:** 10.1101/2025.09.16.676176

**Authors:** Joseph L. Tsenum

**Affiliations:** Phystech School of Biological and Medical Physics, Moscow Institute of Physics and Technology (MIPT), Moscow Region, Russia

**Keywords:** *Long* non-coding RNAs (lncRNAs), DNA sites, triplex structures, deep learning architectures

## Abstract

Long non-coding RNAs (lncRNAs) can perform their regulatory roles by forming triple helices through RNA–DNA interactions. Although this has been verified by a few in vivo and in vitro methods, robust *in silico* approaches that predict the potential of lncRNAs and DNA sites to form triplex structures are still required. Tools such as Triplexator have predicted vast numbers of lncRNAs and DNA sites with triplex forming potential, yet there remains a pressing need for advanced computational methods that can refine and extend these predictions. In this study, we developed ten (10) deep neural network models that predict the potential of lncRNAs and DNA sites to form triple helices on a genome-wide scale. To prepare our dataset, we first used Triplexator to screen out lncRNAs and DNA sites with low triplex-forming potential. We then trained different deep learning architectures, including two-layer convolutional neural networks (CNN), residual neural networks (ResNN), long short-term memory recurrent neural networks (LSTM-RNN), and multilayer perceptron (MLP). Among these architectures, our lncRNA_CNN and LSTM3-RNN both achieved a mean AUC of 0.99 for lncRNA features at a kernel size of 32 and a learning rate of 1e-3. For DNA site features, our DNA_CNN achieved the best performance with a mean AUC of 0.98 under the same conditions. In conclusion, we demonstrate that deep neural network architectures can effectively learn sequence features of lncRNAs and DNA to accurately predict RNA:DNA triplex formation potential, providing a scalable *in silico* framework for studying genome-wide triplex biology.

## Introduction

Advances in genome sequencing technologies in the past decade have shown the complexity of genome organization in both eukaryotic and prokaryotic organisms (Kapranov *et al*., 2007; ENCODE Project Consortium, 2004, 2012). These genome sequencing technologies has revealed large amounts of non-coding RNAs (ncRNAs) that were previously coined as ‘junk DNA’ and a small fraction of protein-coding mRNA in the mammalian DNA.

Efforts have been made in the past years in sequencing ncRNAs and quantifying their various regulatory roles (Quinn and Chang, 2016; Bonasio and Shiekhattar, 2014; Batista and Chang, 2013; Lee, 2012). Non coding RNAs (ncRNAs) consists of a broad class of regulatory RNAs; these include small ncRNAs such as small interferingRNAs (siRNAs), PIWI-interacting RNAs, microRNAs (miRNAs), and long noncoding RNAs (lncRNAs),which are made up of more than 200 nucleotide in length (Cech and Steitz, 2014; Kowalczyk *et al*., 2012). Majority of the mammalian non-coding RNAs are highly conserved, suggesting that they perform significant functions in biological processes (Struhl, 2007).

Long non-coding RNAs (lncRNAs) interacts with epigenetic factors to structure the results of important biological roles such as gene transcription regulation, modulation of nuclear organization, pre-mRNA splicing and X inactivation in females. This has improved our knowledge of the interaction between lncRNAs and epigenetic factors of human diseases as biomarkers of genetic diseases and their therapeutic targets (David J. H *et al*., 2018).

Long non-coding RNAs (lncRNAs) are capable of forming RNA-DNA hybrid duplexes and RNA-DNA triplexes structures at specific DNA sequences (Wang and Chang, 2011; Li *et al*., 2016). These triplex-forming lncRNAs are important in biological processes. Both *in-vivo* and *in-vitro* methods have verified the potential of some lncRNAs in forming triplex structures with DNA sequences. Various mathematical, statistical and computational methods have also been applied in predicting lncRNAs that have the potential of forming triplex structures with specific DNA regions. Computationally, classical machine learning and deep neural networks have shown promise in predicting these triplex-forming lncRNAs and DNA sites.

## Materials and methods

### Data Collection & Preprocessing for our Triplex Forming Prediction Model (TriplexFPM) for Sequence-Based Feature

*531* positive dataset for triplex lncRNAs, *36021* negative dataset for lncRNA triplex; *2542* DNA Site triplex positive dataset and *12,735* DNA Site triplex negative data were collected as follow;

### Triplex lncRNA Prediction Positive Dataset for TriplexFPM

We obtained *531 positive dataset for triplex lncRNAs* were collected from the work of Zhang *et al*., 2020. They collected this positive data for triplex lncRNA prediction in two ways; They extracted lncRNAs according to TriplexRNA regions (DNA:RNA triplex forming peaks in RNA) contained in the work of Sentürk *et al*. by considering both Solid Phase Reversible Immobilization-based paramagnetic bead size selection and immunopurification with anti-DNA antibody RNA separation in Hela S3 cell. They then used GENCODE release 33 lncRNA annotation to extract the lncRNAs that covered the TriplexRNA regions and *476* unique samples were gotten. These lncRNAs were named as triplexlncRNA.

They also obtained a total number of *159* reported triplex lncRNAs that were verified by either *in vivo* or *in vitro* assays to form triplexes with DNA. They obtained these from peer-reviewed papers. These lncRNAs were named reported triplex lncRNA, which included MEG3 (Mondal *et al*., 2015), PARTICL (O’Leary *et al*., 2015), MIR100HG (Wang *et al*., 2018), FENDRR (Navarro *et al*., 2016), and HOTAIR (Kalwa *et al*., 2016). They also took into account all the variants of the reported triplex lncRNA. This brought the number of the collected positive data for triplex lncRNA samples to *635*.Triplex Domain Finder-TDF (Kuo *et al*., 2019) was then used to filter the data since the task is to predict the most likely triplex-forming lncRNAs. In the course of evaluating the triplex forming potential of the *635* lncRNAs with the whole gene promoters (except for chromosome Y and M) with TDF (default parameters), they found out that *104* of them did not contain DNA Binding Domains (DBDs)with powerful Triplex Forming Oligonucleotide (TFO) support.

The challenge of low triplex forming potential gotten from TDF in the collected positive data may be because; for reported triplex lncRNA, each lncRNA gene may have many transcripts with splice variants, but perhaps not all of them have the potential of forming triple helix with DNA, and secondly, for triplexlncRNA, they could not confirm that the overlap regions between R-Loop and TriplexRNA regions as having triplex forming potentials or not (Engreitz *et al*., 2013). To have confidence in the positive dataset, they removed the *104* lncRNAs having low triplex forming potentials and finally, *531* positive triplexlncRNAs dataset were left and we used same in our work.

### Triplex lncRNA Prediction Negative Dataset for TriplexFPM

We obtained *36021 negative datasets for lncRNA triplex* from the work of Zhang *et al*., 2020. They collected the negative lncRNA triplex samples the same way they did for the positive dataset. After removing the lncRNAs in the original positive dataset from GENCODE annotation, they then evaluated the triplex forming potential of all remaining lncRNAs with the whole gene promoters (except for chrY and chrM) using Triplex Domain Finder (TDF). The lncRNAs with at least one powerful Triplex forming oligonucleotides (TFO) supported DBD and at least *123* DNA Binding Sites (DBSs) were kept (the smallest number of DBSs was *123* in positive data). Again, from the screened lncRNAs, they removed *one* lncRNA with letter ‘N’ (variants of lncRNA MALAT1) which was previously confirmed to form RNA-RNA triplex (Ageeli *et al*., 2019). At last, they got *36,021* negative dataset for the lncRNAs that had the comparable potentials of forming triplehelix as their lncRNA negative data and we used same in our task.

### Triplex DNA Sites Prediction Positive Dataset for TriplexFPM

We collected *2542 DNA triplex positive dataset* from the work of Zhang *et al*., 2020.

To predict triplex-forming sites in DNA based on the experimental data, they adopted TriplexDNA regions (DNA:RNA triplex forming peaks in DNA) from the work of Sentürk *et a*l., 2019 as their positive data and we did same. They enriched the RNA-associated DNAs by an unbiased approach. After thy removed *5* samples containing the letter ‘N’ in their sequences, they finally got *2542* DNApositive dataset and we used same in our work.

### Triplex DNA Sites Prediction Negative Dataset for TriplexFPM

They obtained *12,735 DNA triplex negative data* as random regions in promoters. They downloaded all ensembl annotated promoters at Hunt *et al*., 2018, from which *12,735* regions (about 5 times amount of originally positive data) was obtained. They got these regions by randomly selecting chromosomes (except chrY and chrM) and DNA regions with the same lengths as TriplexDNA. They extracted the sequence data for both positive and negative from the DNA minus strand.

**Fig 1.1:**
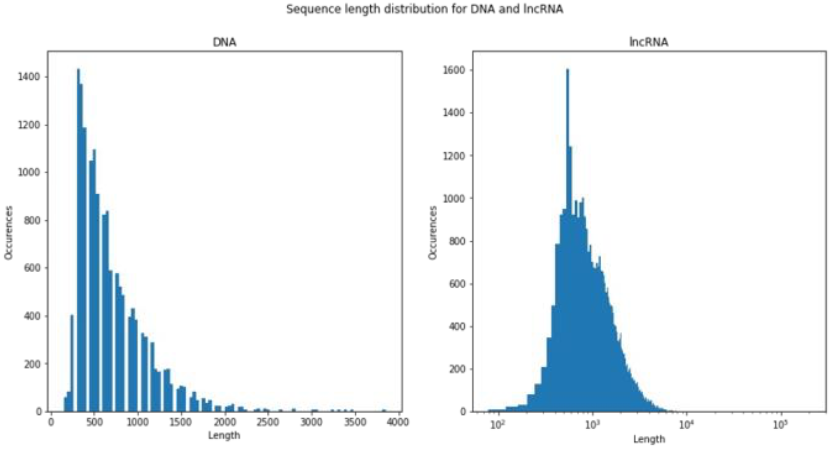
Triplex lncRNA and DNA site length distributions for TriplexFPM

### Model Construction for TriplexFPM

We imported the libraries we needed from python (mainly Tensorflow and Keras), biopython (mainly SeqIO) and others. We then denoted some supplementary functions that were needed to extract sequence features used for training/testing. We balanced our data by enriching our positive data by oversampling and depleting our negative data by undersampling. We fixed the data size problem for lncRNA triplex as lncRNA_samples = 1500 (3 times bigger than the size of the positive samples), while the data size for DNA triplex as DNA_samples = 6000 (2.4 times biggerthan the size of the positive samples).

### Feature Extraction

K-mer is a commonly used approach to transform a sequence into a vector. It usually counts the frequencies of single or multiple nucleotide compositions in a sequence and represents the sequence into a 4 k dimensional vector while kmerscore is an overall measure of the k-mer nucleotide composition bias in a sequence, it is extracted from k-mer features (Zhang *et al*., 2020). While for our CNN and ResNN, in order to extract our features, we used *K-mer and kmerscore from 1-6*, for our LSTM RNN, we used one-hot encode.

After preprocessing our data, we merged the positive and negative data, created label arrays, shuffled our data and saved our sequence data, labels and features as *processed data* for model construction. We then defined our constant hyperparameters by splitting our data into *train, validation and test* datasets with *80%, 10% and 10%* respectively as shown below:

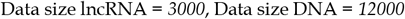

We reloaded the processed_data files and scaled features for both datasets using standard scaler to enable the model to calculate distances between data for better model performance. Standardization transforms the data to have zero mean and a variance of 1, they make our data unitless (Baijayanta, 2020). Taking lncRNA and DNA sites as our features, we adopted a 2-layerConvolutional Neural Network (CNN) to construct our CNN models, which effectively learned high-level features from our data. We also used LSTM-RNN and ResNN to train our models. We first trained *5 models* predict the most likely triplex-forming lncRNAs in practice from the lncRNAs triplex forming potentials data as estimated by computational methods. We also trained *5* other models to predict the potential of DNA sites in forming triplex structures from the experimentally verified data as reported by Zhang *et al*., 2020.

**Fig 1.2:**
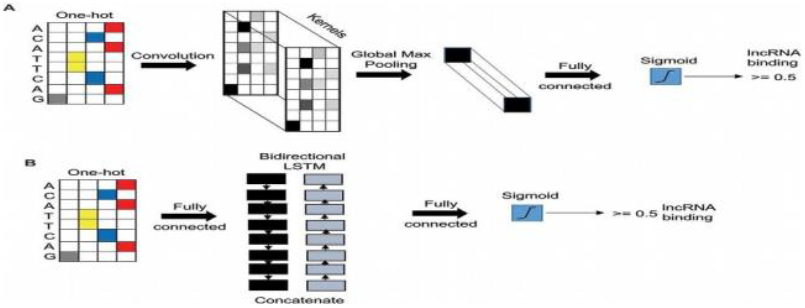
Ideas for Model construction for TriplexFPM (Wang *et al*., 2020)

### Hyperparameter Tuning for TriplexFPM

We defined our internal and external hyperparameters for our CNN, and ResNN as follows: epochs (50), input shape of (90, 1), learning rate at 1e-3 for 32 Kernel size (which performed better) and we set the verbose at 1 just to see how the training progress for each epoch, ReLU as our hidden layer activation function and Sigmoid as our output activation function. For the LSTM RNN models, our internal and external hyperparameters were as follow: epochs (80), input shape of (none, 4), learning rate at 1e-3 for 32 Kernel size, ReLU as our hidden layer activation function and Sigmoid as our output activation function. We used Adam optimization algorithm as our optimizer and since it is a binary classification task, we used binary cross entropy as our loss function. We used six different metrices to assess the performance of our models in all the architectures, including: binary accuracy, precision, recall and AUC, F1-score, MCC. We employed early stopping criteria on the optimization procedure when the values of the loss function on the validation set stopped decreasing at a particular consecutive time.

## Results and discussion

We trained a total of ten (10) deep neural network models based on lncRNA and DNA site features. These models constitute our Triplex Forming Prediction Model (TriplexFPM) and include:

∘ **lncRNA-based models:** lncRNA_CNN, lncRNA_ResNN, lncRNA_LSTM1-RNN, lncRNA_LSTM2-RNN, lncRNA_LSTM3-RNN.
∘ **DNA-based models:** DNA_CNN, DNA_ResNN, DNA_LSTM1-RNN, DNA_LSTM2-RNN, DNA_LSTM3-RNN.

### Performance of TriplexFPM (lncRNA/DNA Models)

Our feature-based models trained on lncRNA and DNA sequences (TriplexFPM) demonstrated consistently high predictive performance across multiple architectures (Table 1.1). Among the lncRNA-based models, **lncRNA_CNN** and **lncRNA_LSTM3-RNN** achieved the highest performance with a mean AUC of **0.99** at a kernel size of 32 and a learning rate of 1e-3, indicating that convolutional and recurrent architectures are both capable of effectively capturing lncRNA sequence patterns relevant to triplex formation. This was closely followed by **lncRNA_LSTM2-RNN** (AUC = 0.98) and **lncRNA_ResNN** (AUC = 0.97), while **lncRNA_LSTM1-RNN** performed lowest (AUC = 0.92).

**Table 1.1:**
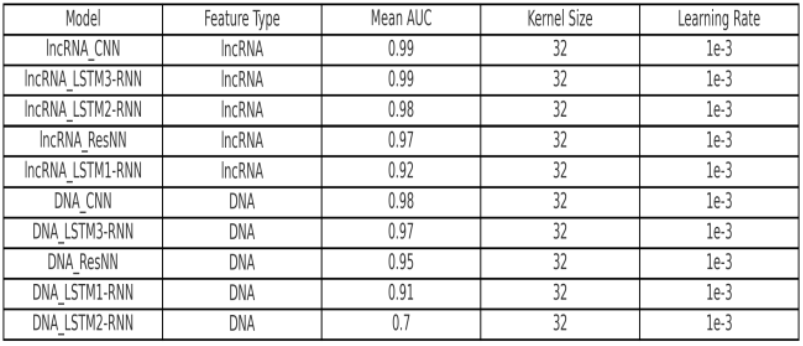
TriplexFPM (lncRNA/DNA feature-based models)

**Fig 2.1:**
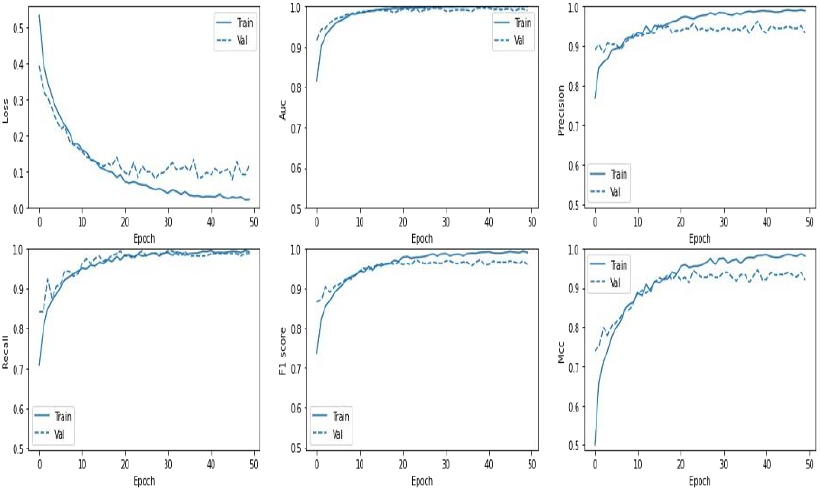
CNN Performance Plot for lncRNA TriplexFPM

**Fig 2.2:**
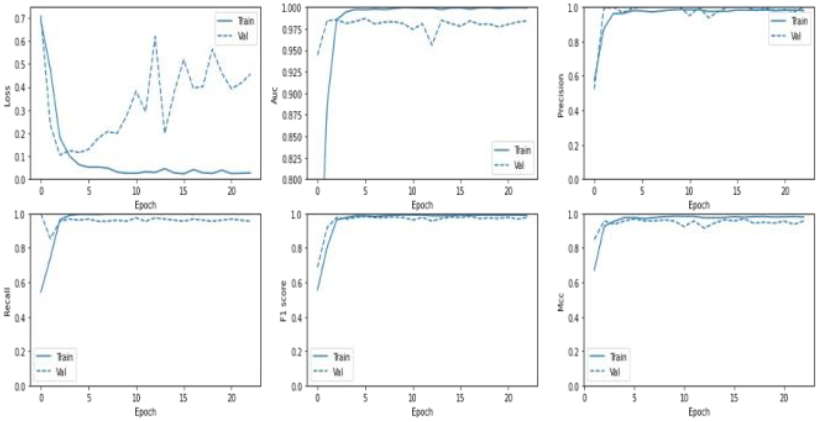
LSTM3-RNN performance plot for lncRNA TriplexFPM

For DNA feature-based models, **DNA_CNN** exhibited the best performance with a mean AUC of **0.98**, followed by **DNA_LSTM3-RNN** (AUC = 0.97) and **DNA_ResNN** (AUC = 0.95). The relatively weaker performance of **DNA_LSTM1-RNN** (AUC = 0.91) and **DNA_LSTM2-RNN** (AUC = 0.70) suggests that certain recurrent architectures may be less effective at capturing DNA sequence signals compared to convolutional and residual networks.

**Fig 2.3:**
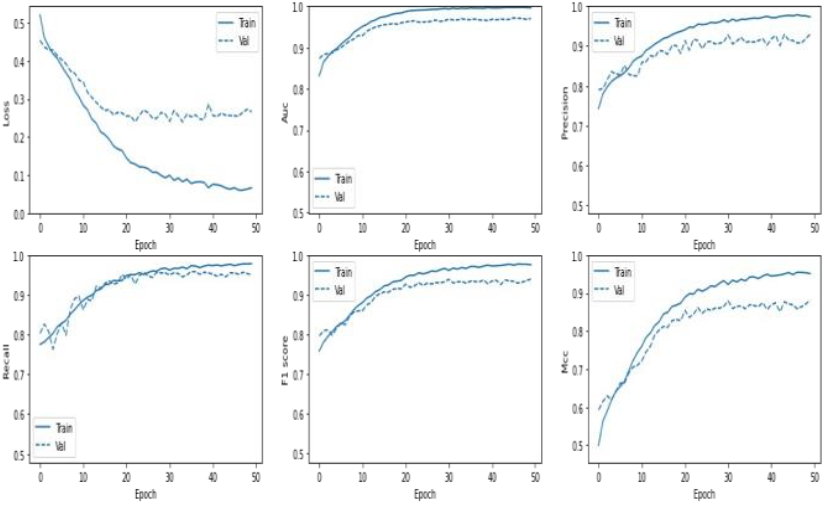
CNN performance plot for DNA site TriplexFPM

**Fig 2.4:**
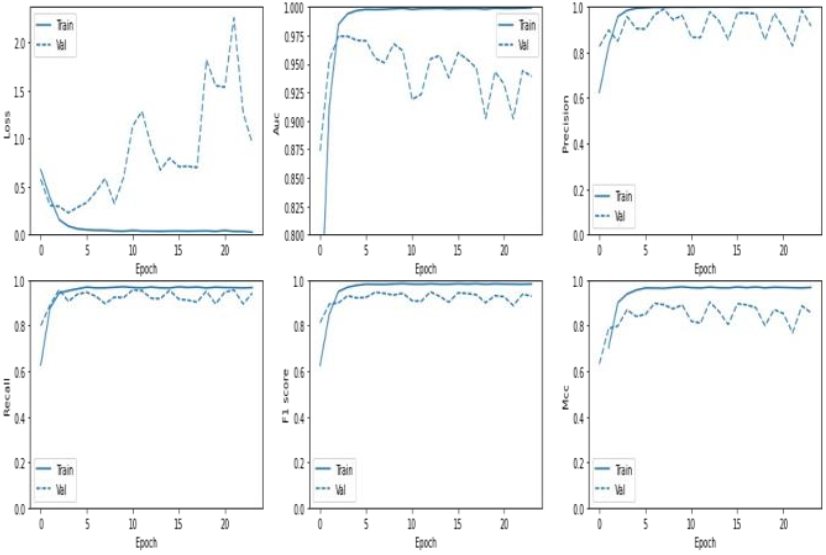
LSTM3 RNN performance plot for DNA site TriplexFP

**Fig 2.5:**
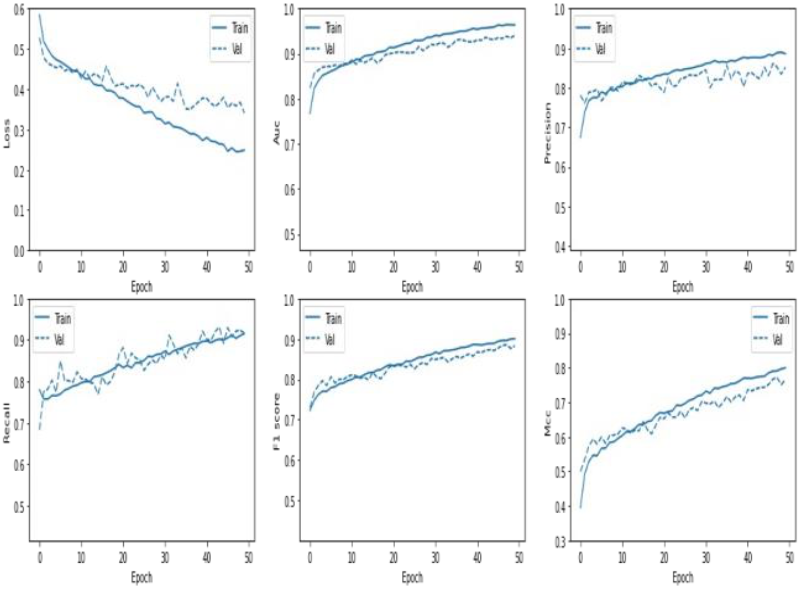
ResNN performance plot for DNA site TriplexFPM

Overall, the high area under the ROC curve (AUC) values across most architectures indicate that the lncRNAs and DNA sites identified by Triplexator as having triplex-forming potential indeed contain strong sequence signals that can be captured by deep neural networks. The superior performance of convolutional models highlights the importance of localized motif detection in triplex prediction, while the variability among recurrent architectures suggests the need for careful tuning of sequence-dependent models.

**Fig 2.6:**
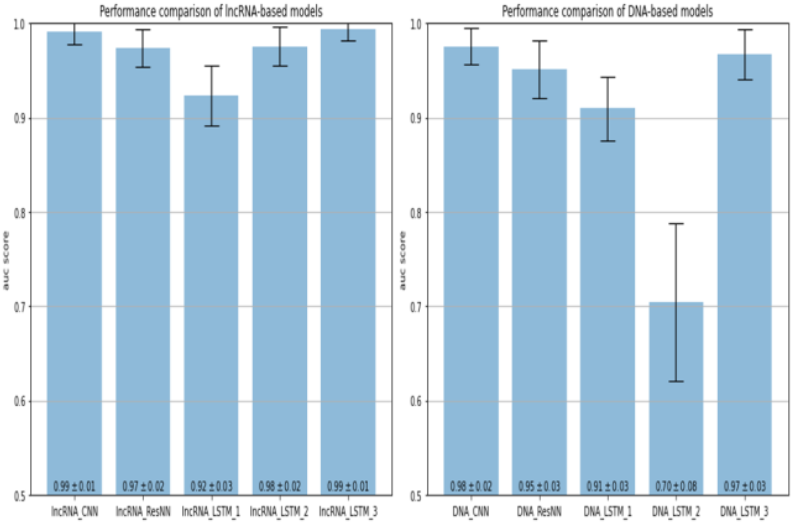
Performance comparison of different lncRNA & DNA site feature-based models on mean AUC

### Limitations

One of the challenges encountered in this study was the limited availability of lncRNA positive datasets with confirmed triplex-forming potential. Deep learning models generally benefit from large and diverse datasets, and the relatively small positive dataset used here may have constrained the performance of some models across specific metrics. In addition, since our dataset was collected from previously published studies and public repositories rather than generated experimentally, discrepancies in data quality and annotation could have contributed to inconsistencies in performance and reduced confidence in some results.

### Future Directions

To address these challenges, future work should expand training beyond promoter regions to include additional genomic contexts, incorporating histone modification marks as features to further refine triplex prediction. Training different DNA regions separately and later integrating them may also improve performance. Beyond deep learning, classical machine learning models such as support vector machines, random forests, and decision trees should be benchmarked against our models to evaluate their comparative effectiveness. From a computational standpoint, implementing models in PyTorch instead of Keras-TensorFlow may provide greater flexibility and optimization opportunities. Importantly, optimizing kernel sizes and learning rates for each performance metric rather than applying uniform hyperparameters may yield further improvements. Finally, experimental validation of the predicted triplex-forming lncRNAs and DNA sites using in vitro and in vivo assays remains essential. Such efforts could establish biomarkers for complex human diseases and open new avenues for therapeutic intervention.

## Declarations

### Ethics Approval and Consent to Participate

Not applicable

### Consent for Publication

Not applicable

### Availability of Data and Materials

The datasets and deep learning models used in this study are openly available on GitHub at https://github.com/Joseph-Luper-Tsenum/Predicting-RNA-DNA-Triplex-Structures-from-Sequence-Features-Using-Deep-Learning-Architectures/tree/main

### Authors’ Contribution

JLT designed the study, performed the analyses, and wrote the manuscript.

## Acknowledgments

The authors thank the Open Doors Olympiad for providing the scholarship, the Russian Government for funding, Phystech School of Biological and Medical Physics (FBMP) at the Moscow Institute of Physics and Technology (MIPT) for their support, and the members of the International Laboratory of Bioinformatics at HSE University for their valuable assistance and guidance.

## Abbreviations

CNN: Convolutional neural network
RNN: Recurrent neural network
ResNN: Residual neural network
LSTM: Long short-term memory
MLP: Multilayer perceptron
RNA: Ribonucleic acid
DNA: Deoxyribonucleic acid
lncRNA: long non-coding RNA
TriplexFPM: Triplex Forming Prediction Model
AUC: Area under the Receiver operating characteristic (ROC) Curve
ReLU: Rectified Linear Unit
lr: Learning rate
MCC: Matthews correlation coefficient
k-mer: A k-mer is a subsequence of length k derived from a longer nucleotide or amino acid sequence

## Supplementary Figures

Additional results of the **TriplexFPM** model are provided in the supplementary figures and tables. These include extended performance metrics, comparative analyses, and visualizations that complement the main text.

### A. lncRNA-based models: lncRNA_ResNN, lncRNA_LSTM1-RNN, lncRNA_LSTM2-RNN

**Fig 3.1:**
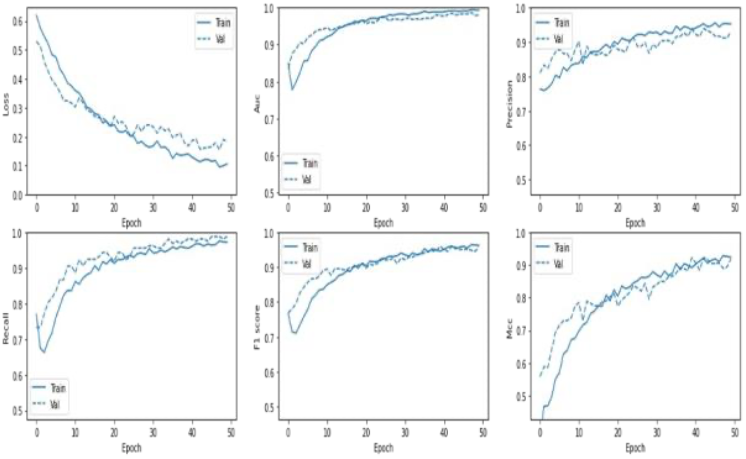
ResNN performance plot for lncRNA TriplexFPM

**Fig 3.2:**
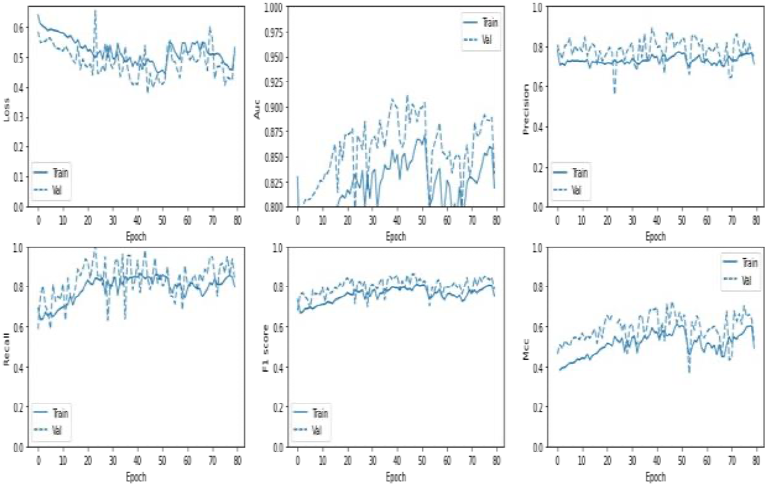
LSTM1-RNN performance plot for lncRNA TriplexFPM

**Fig 3.3:**
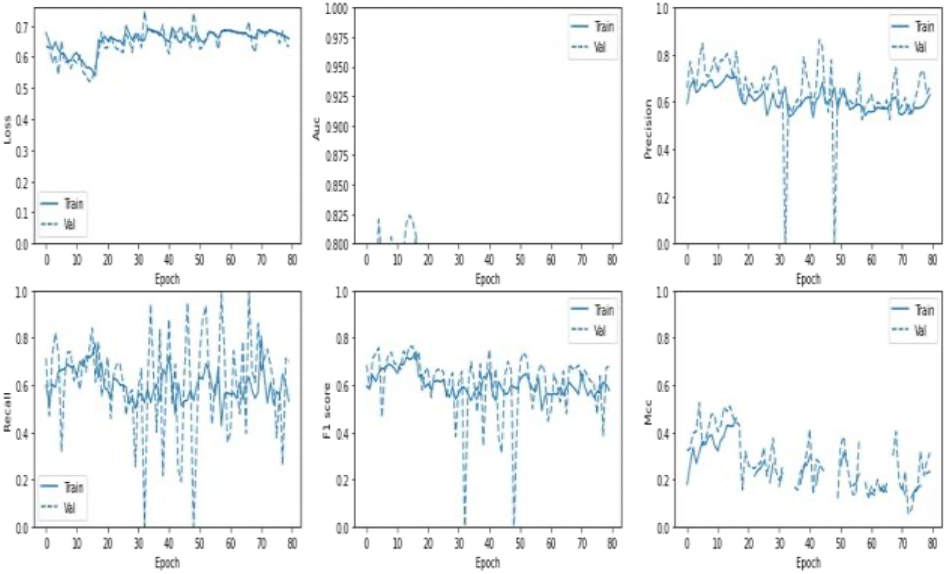
LSTM2-RNN performance plot for lncRNA TriplexFPM

### B. DNA-based models: DNA_ResNN, DNA_LSTM1-RNN, DNA_LSTM2-RNN

**Fig 4.1:**
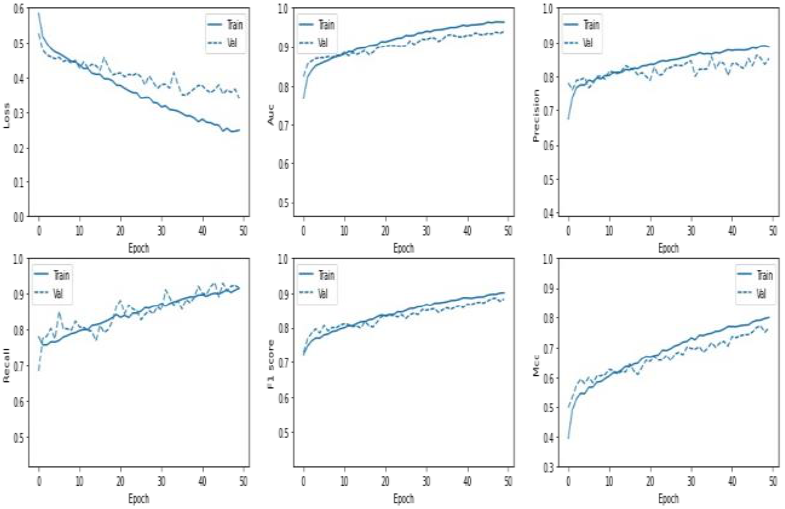
ResNN performance plot for DNA site TriplexFPM

**Fig 4.2:**
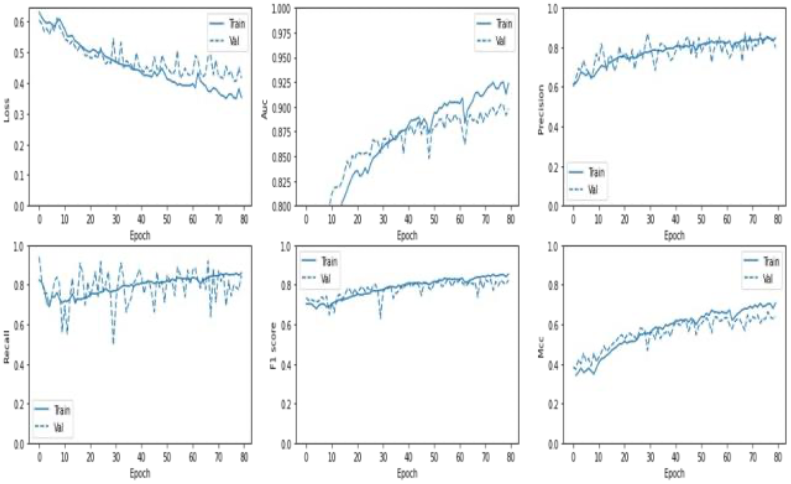
LSTM1-RNN performance plot for DNA site TriplexFPM

**Fig 4.3:**
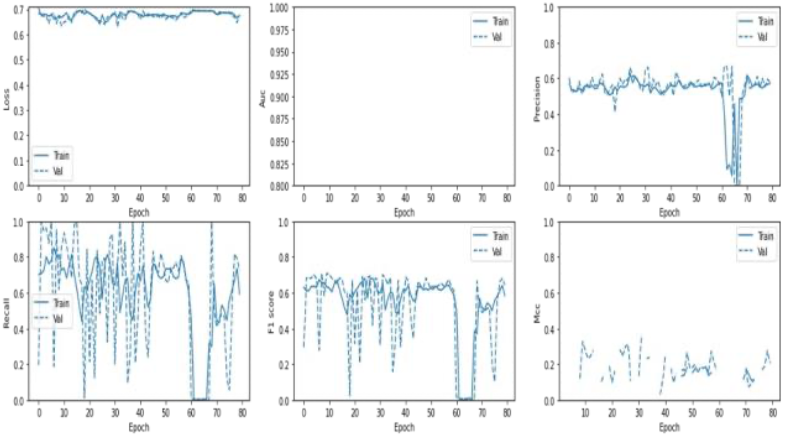
LSTM2-RNN performance plot for DNA site TriplexFPM

### C. Performance comparison of different lncRNA & DNA site feature-based models

**Fig 5.1:**
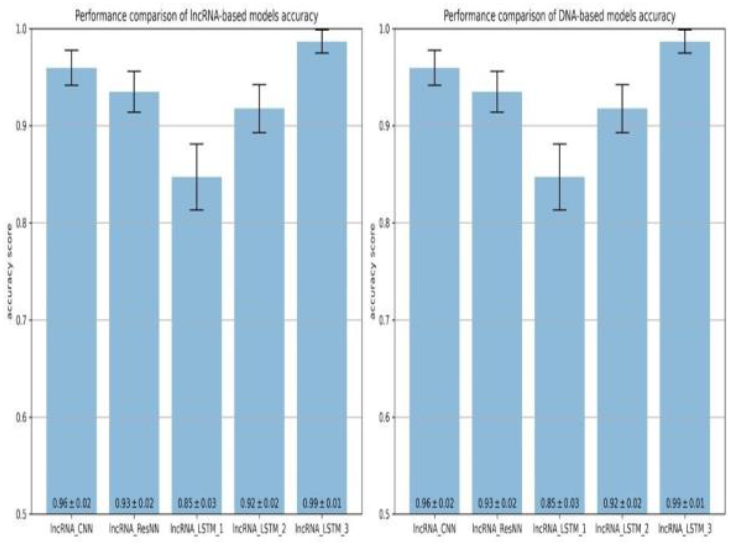
Performance comparison of different lncRNA & DNA site feature-based models on mean accuracy

**Fig 5.2:**
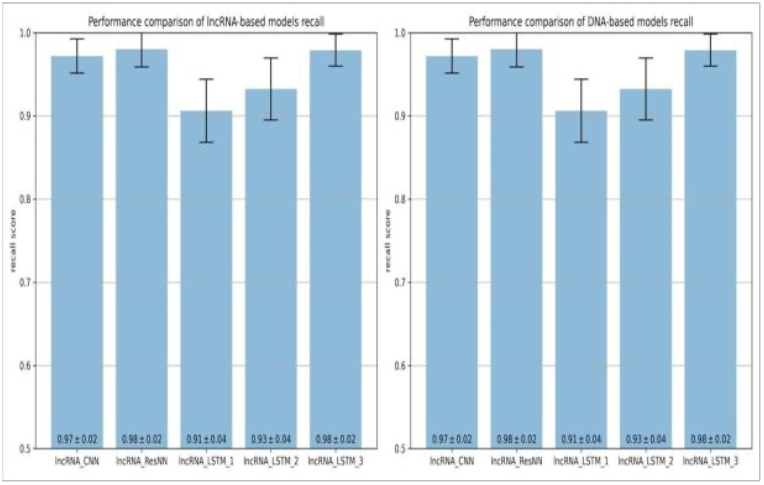
Performance comparison of different lncRNA & DNA site feature-based models on mean recall

**Fig 5.3:**
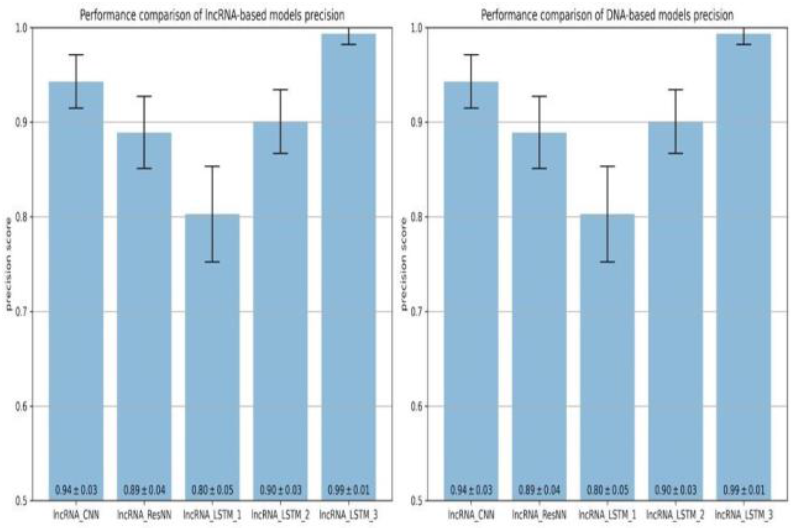
Performance comparison of different lncRNA & DNA site feature-based models on mean precision

**Fig 5.4:**
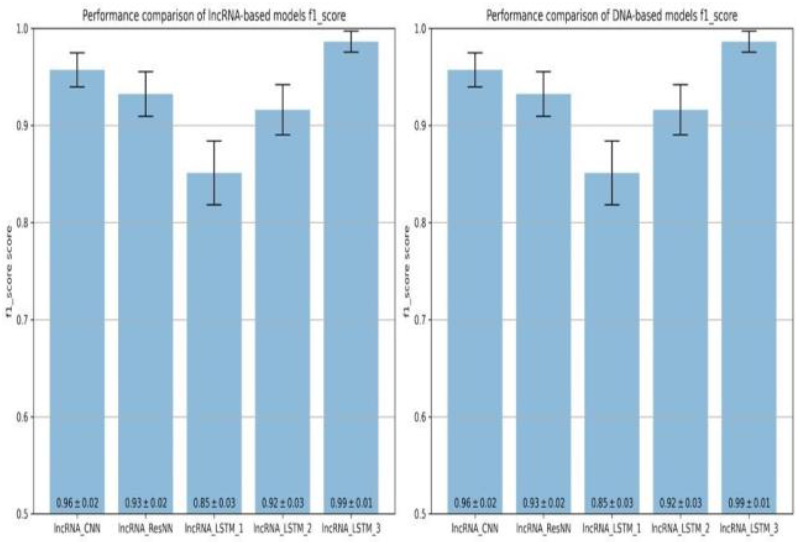
Performance comparison of different lncRNA & DNA site feature-based models on mean F1-score

**Fig 5.5:**
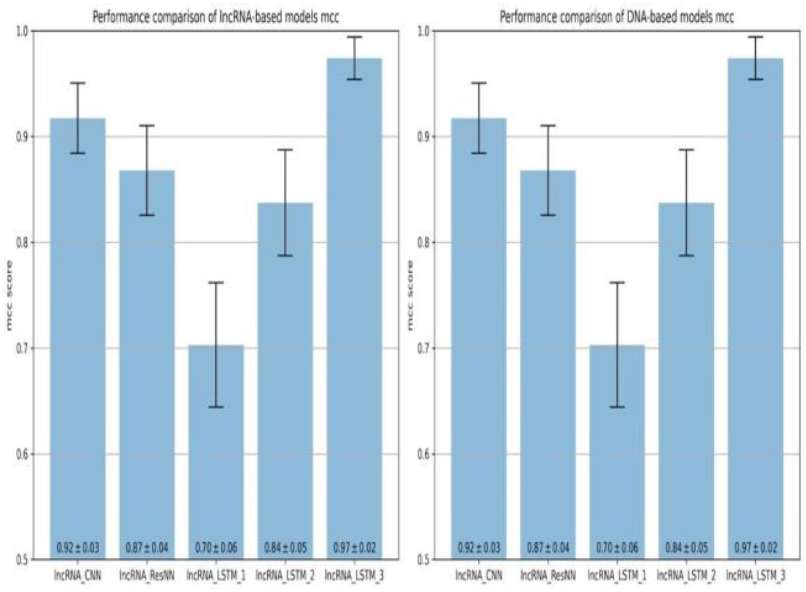
Performance comparison of different lncRNA & DNA site feature-based models on mean MCC

